# Interplay between autophagy, the unfolded protein response and the cytoplasmic heat stress response during heat stress in maize

**DOI:** 10.1101/2024.11.06.622333

**Authors:** Jie Tang, Zhaoxia Li, Diane C Bassham

## Abstract

High temperatures can substantially reduce plant survival and reproduction, and therefore decrease crop productivity. Plants activate several pathways in response to high temperatures, including the endoplasmic reticulum (ER) unfolded protein response (UPR), the cytoplasmic heat stress response, and autophagy, which together aid in stress tolerance by refolding or degrading misfolded and unfolded proteins. The relationship between the UPR and autophagy is known, as ER stress activates the UPR, and a key activator of the UPR, Inositol-requiring enzyme type 1 (IRE1B), upregulates autophagy. To assess the relationship between the distinct responses to heat stress in maize, we analyzed the effect of disruption of the UPR, via a *bzip60* mutant, on autophagy. We found that genes related to autophagy are upregulated in the *bzip60* mutant, and that autophagy is activated in this mutant even in the absence of stress. We in turn analyzed the effect of loss of autophagy on the UPR. The *bZIP60* mRNA spliced form, a marker for the UPR, was increased in mutants in *ATG10*, a core component of the autophagy machinery, during ER stress. By contrast, *bZIP60* splicing was unaffected in *atg10* mutants in response to heat, whereas cytoplasmic heat stress components were increased. Loss of autophagy therefore differentially affects other heat stress response pathways, depending on the specific stress conditions leading to the activation of the pathways.

## Introduction

Extreme temperatures have severe effects on agricultural production, damaging food security (Lobell *et al*., 2011; Lesk *et al*., 2016), and the cellular and physiological responses of plants to these conditions are vital for survival (Zhang *et al*., 2018). Understanding the mechanisms by which plants cope with heat stress is critical in order to produce plants with increased thermotolerance (Ohama *et al*., 2017). These mechanisms include the production of heat shock proteins (HSPs) through a cytoplasmic heat stress response, the unfolded protein response (UPR), together with ER-associated degradation (Chen *et al*., 2020), within the endoplasmic reticulum (ER), and activation of protein and organelle degradation via autophagy. How these processes are appropriately coordinated, and how the signaling pathways leading to their activation are integrated, is unclear (Li and Howell, 2021).

Autophagy, literally defined as “self-eating”, is highly conserved among eukaryotes and is critical for stress tolerance. In plants, it is activated to recycle cytoplasmic contents during development and in response to stress, and a basal level of autophagy is important for cellular homeostasis (Wang *et al*., 2018) and in balancing growth with stress tolerance (Avin-Wittenberg *et al*., 2018; Tang and Bassham, 2018).

Upon activation of autophagy, a double-membrane autophagosome encloses cargo and transports it to the vacuole for degradation (Yang and Bassham, 2015; Soto-Burgos *et al*., 2018). The identification of autophagy-related (ATG) genes in yeast (Tsukada and Ohsumi, 1993; Harding *et al*., 1995; Thumm *et al*., 1994) was key in understanding the mechanism by which autophagy occurs. The core machinery for autophagosome formation includes the autophagy-initiating ATG1 kinase complex (Kamada *et al*., 2000); two ubiquitin-like conjugates, ATG12-ATG5 and ATG8-PE, which localize to the phagophore assembly site (PAS) and function in autophagosome formation (Le Bars *et al*., 2014; Yin *et al*., 2016); and ATG9/ATG2/ATG18, which may function in expansion of the forming autophagosome from the ER via lipid transport activity (Zhuang *et al*., 2017; Gomez-Sanchez *et al*., 2018; Valverde *et al*., 2019; Matoba *et al*., 2020; Aroca *et al*., 2021). In plants, autophagy is activated in response to stress conditions, including nutrient deficiency (Doelling *et al*., 2002; Hanaoka *et al*., 2002), abiotic (Avin-Wittenberg, 2019) and biotic stress (Liu *et al*., 2005; Lai *et al*., 2011). Arabidopsis mutants in *ATG* genes are hypersensitive to numerous environmental stresses, illustrating the key role of autophagy in plant stress resilience (Avin-Wittenberg, 2019).

When plants encounter elevated temperatures, several response pathways are activated to allow tolerance of these stress conditions. Major pathways for heat tolerance include the cytoplasmic heat stress response (HSR) and the UPR in the ER (Li and Howell, 2021).

Errors in protein folding within the ER upon stresses such as high temperatures lead to the accumulation of misfolded proteins, a potentially toxic condition termed ER stress. ER stress induces adaptive responses including the UPR and autophagy (Liu *et al*., 2012), which mitigate the damage caused by stress and protect plants from further stress. ER stress-induced autophagy helps to degrade pieces of the ER containing misfolded proteins (Yang *et al*., 2016). In Arabidopsis and maize, autophagy can be activated by the ER stress agents tunicamycin and dithiothreitol (DTT), both of which cause the accumulation of unfolded proteins (Liu *et al*., 2012; Srivastava *et al*., 2018), and potentially mimic the effects of heat on the UPR. Plants have two branches in the signaling pathways leading to activation of the UPR, and the UPR signaling arm involving the non-conventional splicing factor IRE1b plays a critical role in the induction of autophagy in response to ER stress, but not to other stress conditions (Liu *et al*., 2012; Li and Howell, 2021). However, heat stress-induced autophagy is only partially suppressed in an *ire1b* mutant (Yang *et al*., 2016), suggesting that other mechanisms for activating autophagy are also important during heat stress.

The misfolding of proteins brought about by heat stress can also lead to protein aggregation within the cytoplasm. This triggers the cytoplasmic HSR, in which heat shock transcription factors (HSFs) activate the expression of HSPs, molecular chaperones that prevent protein misfolding and aggregation. A major class of HSFs are sequestered in the cytoplasm in nonstress conditions and are released to enter the nucleus upon heat stress, where they bind to heat shock response elements in the promoters of HSPs and activate their expression (Li and Howell, 2021). Transcriptomic, metabolomic and proteomic analyses in maize after heat stress (Frey *et al*., 2015; Shi *et al*., 2017; Zhang *et al*., 2020; Abou-Deif *et al*., 2019) have demonstrated the dramatic effects of heat on cell metabolism and the coordination of response pathways.

Under severe or extended heat stress, the HSR may be insufficient to prevent protein aggregation in the cytoplasm. Protein aggregates are potentially toxic, and can be selectively targeted for degradation by a form of autophagy called aggrephagy (Øverbye *et al*., 2007). Aggrephagy selects protein aggregates for degradation via binding to selective autophagy receptors, which recruits them to forming autophagosomes (Luong *et al*., 2022; Clavel and Dagdas, 2021). In tomato, heat shock tolerance requires an active autophagy pathway (Zhou *et al*., 2014), and the upregulation of autophagy genes in response to heat requires the transcription factor HSFA1a (Wang *et al*., 2015). One function of autophagy during heat stress may be to clear cytoplasmic protein aggregates, thus preventing their cytotoxicity (Jung *et al*., 2020).

It therefore appears that in plants, autophagy activation is coordinated with both the cytoplasmic HSR and the ER UPR. Crosstalk between the HSR and the UPR have been demonstrated in tomato, in which overexpression of HSFA1a increases the activation of the UPR, and also upregulates the expression of several autophagy-related genes (Löchli *et al*., 2023). In maize, we have shown that the activity of the *HSF13* promoter is regulated by bZIP60, and a *bzip60* mutant has decreased expression of cytoplasmic HSR genes (Li *et al*., 2020). These results indicate a complex relationship between the ER UPR, cytoplasmic HSR and autophagy that warrants further investigation.

We have shown previously that maize autophagy is activated both by ER stress agents and by heat, both in controlled environment chambers and in the field (Li *et al*., 2020), although the relationship with other heat stress responses is not yet clear. Here, we extend these findings to show that autophagy is constitutively upregulated in a *bzip60* mutant, and that the UPR is upregulated in an autophagy-defective mutant in response to ER stress, whereas the HSR is upregulated in response to heat in the same mutant. The loss of autophagy in maize therefore leads to distinct responses depending on the specific stress encountered.

## Results

### Autophagy is activated by ER stress agent DTT in maize

We have shown previously that autophagy is activated by tunicamycin in roots of maize germinating seeds, and upon exposure to high temperatures (Srivastava *et al*., 2018; Li *et al*., 2020). As in Arabidopsis, autophagy triggered by heat stress is dependent on the ER stress response factor IRE1b (Liu *et al*., 2012), we hypothesized that autophagy during maize heat stress is due to the accumulation of unfolded proteins in the ER, leading to activation of ER stress responses.

We first established the effect of ER stress in 3-leaf stage seedlings of maize by treating seedlings with DTT. DTT is a redox reagent that can disrupt protein folding by affecting the formation of disulfide bonds (Howell, 2013) and has been extensively used as both an ER stress inducer and autophagy inducer in Arabidopsis (Deng *et al*., 2011; Liu *et al*., 2012; Yang *et al*., 2016). Maize W22 seeds were planted in germination papers and grown vertically in water for 5 days, and then treated with or without 2 mM DTT for 6 days. As ER stress has been shown to inhibit plant growth in Arabidopsis (Yang *et al*., 2016), we measured the fresh weight of shoots and roots of seedlings with or without DTT treatment. As predicted, DTT treatment caused a significant reduction in the fresh weight of both shoots and roots (**Figure 1**).

**Figure 1.**
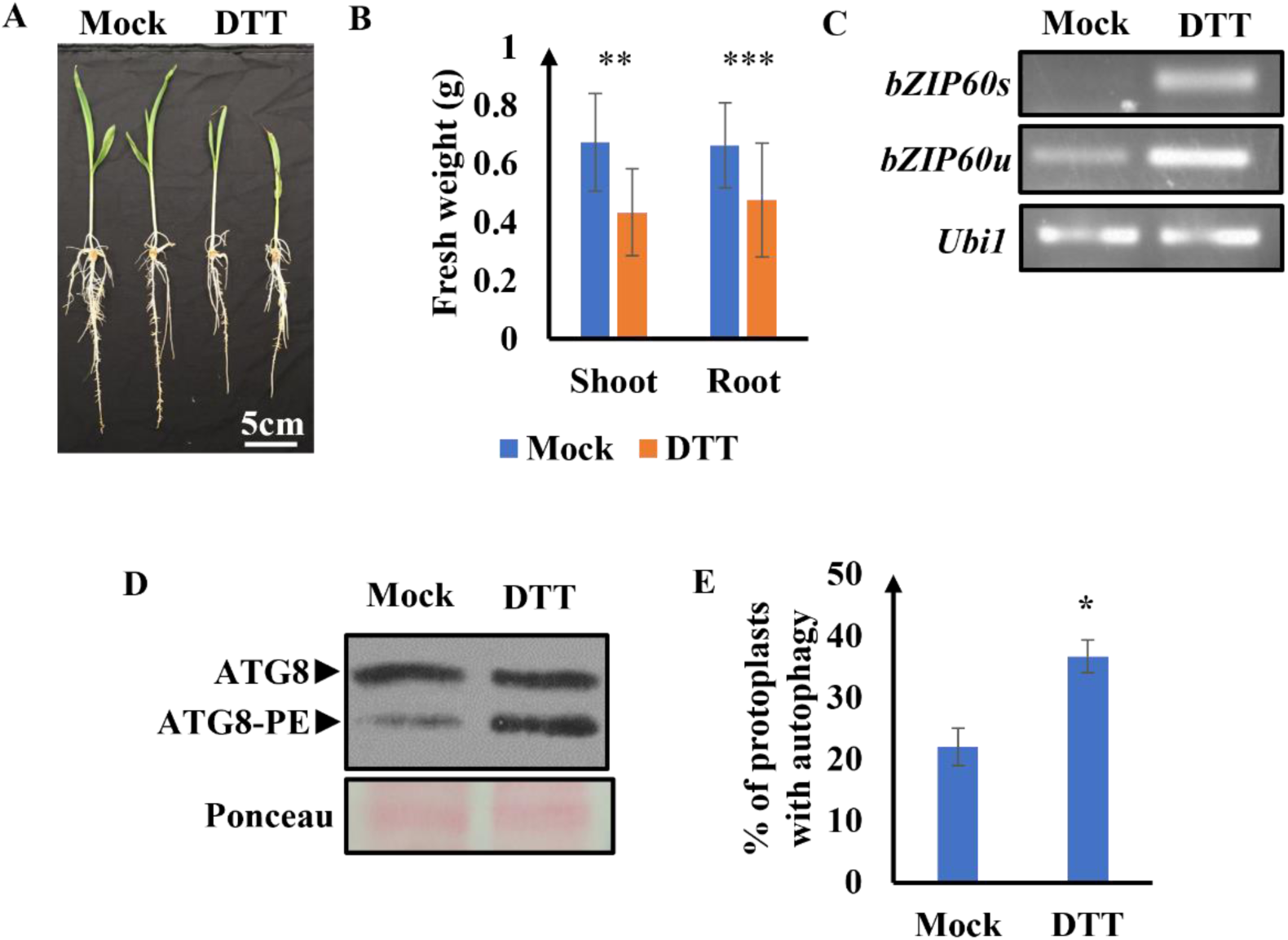
DTT triggers autophagy in maize seedlings. Maize W22 seeds were planted in germination papers and grown vertically in water for 5 days, followed by treatment with or without 2 mM DTT for 6 days. **A**. A representative image comparing the growth of mock-treated seedlings and DTT-treated seedlings. **B**. Quantification of shoot and root fresh weight after mock or DTT treatment. Asterisks indicate statistically significant differences according to Student’s t test (**p<0.01, ***p<0.001). **C**. Activation of the UPR was demonstrated by measuring *bZIP60* mRNA splicing. cDNA was synthesized from 1 µg total RNA for each treatment, followed by PCR using specific primers to amplify unspliced *bZIP60* and spliced *bZIP60*. *Ubiquitin1* (*Ubi1*) was used as a control. **D**. Autophagy activity was measured using an ATG8 lipidation assay. 20 µg total root protein for each treatment was separated by SDS-PAGE using 15% polyacrylamide gels with 6 M urea in the resolving gel, and analyzed by immunoblotting with anti-ATG8 antibody. Ponceau staining was used to indicate equal loading. A representative picture was shown; three biological replicates showed similar results. **E**. Activation of autophagy by ER stress in leaf mesophyll protoplasts. Protoplasts were isolated from soil-grown 2-week-old wild-type W22 seedlings and transformed with GFP-ATG8e. After overnight incubation, protoplasts were treated with water as mock or 2 mM DTT for 6 h. Autophagosomes were visualized and quantified using fluorescence microscopy, and the percentage of protoplasts with activated autophagy was calculated. Asterisk indicates statistical significance, p<0.05. Error bars represent means±SD for 3 biological replicates.

To determine the extent of activation of the UPR upon DTT treatment, splicing of *bZIP60* mRNA was measured as a marker for the UPR (Deng *et al*., 2011; Li *et al*., 2020). The spliced form and the unspliced form of bZIP60 both increased in DTT-treated roots when compared to mock-treated roots (**Figure 1C**), indicating that the UPR is activated. To test if autophagy is also activated by DTT, proteins were extracted from roots and ATG8 lipidation analyzed as described previously (Srivastava *et al*., 2018; Li *et al*., 2020). Increased ATG8-PE was observed in DTT-treated roots compared to mock-treated roots (**Figure 1D**), suggesting that autophagy is induced. To confirm the induction of autophagy by DTT, we transiently expressed the fluorescent autophagosome marker GFP-ZmATG8e in leaf protoplasts isolated from 8- to 10-day-old W22 seedlings. Since ATG8 is localized on autophagosomes, fluorescent protein tagged ATG8 is commonly used as an effective marker for visualization of autophagy (Yoshimoto *et al*., 2004; Contento *et al*., 2005; Li *et al*., 2015). GFP-ZmATG8e-expressing protoplasts were treated with or without 2 mM DTT for 6 to 8 hours and autophagosomes were quantified by fluorescence microscopy. Protoplasts with more than 3 puncta were considered to have activated autophagy, and the percentage of protoplasts with activated autophagy was calculated. Upon DTT treatment, a significantly higher percentage of showed activated autophagy compared to the mock treatment (**Figure 1E**), further confirming that autophagy can be activated by DTT in maize.

### Autophagy is constitutively activated in a *bzip60-2* mutant

As autophagy is activated by ER stress, we hypothesized that when the UPR is deficient, leading to accumulation of unfolded ER proteins, autophagy may be upregulated to increase protein degradation capacity to compensate. We have recently characterized a maize *bzip60* mutant, *bzip60-2* (mu016844), with greatly decreased expression of *bZIP60* and defective UPR activation (Li *et al*., 2020). The effect of various maximum daily temperatures (31°C, 33°C, 35°C, 37°C) on gene expression in W22 and *bzip60-2* plants was analyzed by RNAseq (Li *et al*., 2020), and loss of bZIP60 was found to affect both UPR and HSR gene expression in response to heat. Several autophagy-related genes were significantly upregulated in the *bzip60-2* mutant when compared to W22 plants, even under non-stress conditions (**Figure 2A**). Thus, we hypothesized that autophagy is upregulated in the *bzip60-2* mutant. To test this hypothesis, maize W22 and *bzip60-2* seeds were planted in rolls of germination paper. Plants were grown in water for 5 days and then treated with or without 2 mM DTT for 6 days. Roots were collected for protein extraction and ATG8 lipidation assessed. Consistent with previous results (**Figure 1D**), in WT samples ATG8-PE was at a low level upon mock treatment and was significantly increased in DTT-treated roots (**Figures 2B and 2C**). By contrast, in *bzip60-2* mutants, ATG8-PE hyperaccumulated even after mock treatment when compared to W22 plants, and was not further increased under DTT treatment (**Figures 2B and 2C**). These results suggest that autophagy is constitutively activated in a *bzip60-2* mutant.

**Figure 2.**
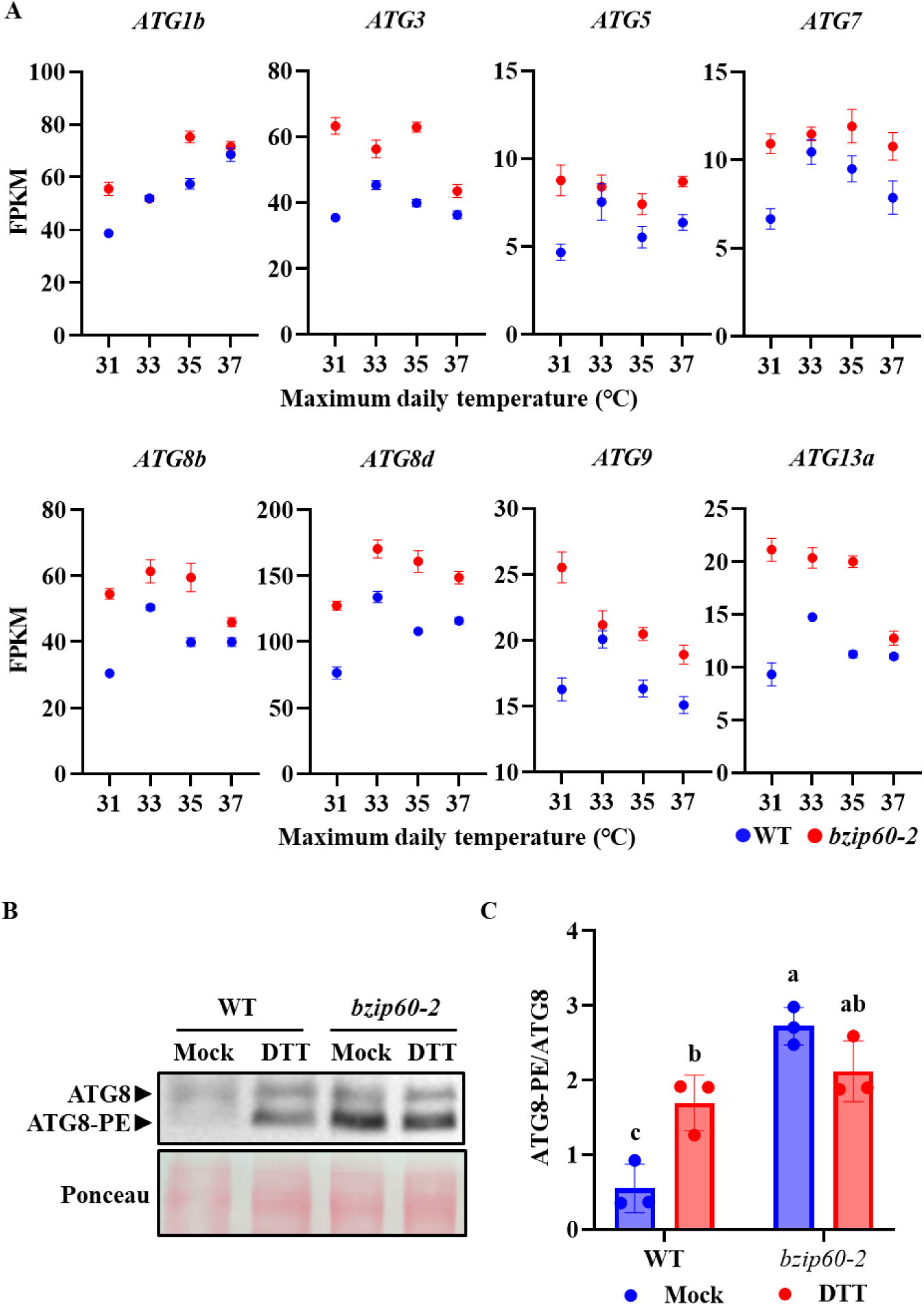
A *bzip60-2* mutant has constitutive autophagy. **A.** Box plots show the relative expression levels (Fragments Per Kilobase of transcript per Million mapped reads) of *ATG* genes in *bzip60-2* and W22 at different maximum daily temperatures at the V5 stage (27 DAG), as determined by RNAseq analysis. **B.** Maize W22, and *bzip60-2* seeds were planted in rolls of germination papers. Plants were grown in water for 5 days and then treated with or without 2 mM DTT for 6 days. Roots were collected for protein extraction and protein separated by SDS-PAGE with 6 M urea in the resolving gel, followed by immunoblot analysis with anti-ATG8 antibody. Ponceau staining was used to indicate equal loading. A representative picture is shown; three biological replicates showed similar results. **C.** Quantification of the ratio of ATG8-PE/ATG8 from Figure 2B. Different letters indicate significant differences, p<0.05, via Tukey’s multiple comparison test.

### Autophagy is deficient in *atg10* mutants

To address the role of autophagy in maize ER stress and heat stress responses, we identified autophagy-deficient mutants from among transposon insertion lines affecting various *ATG* genes from the *UniformMu* (McCarty *et al*., 2005; Settles *et al*., 2007) and the *Ac/Ds* collections (Vollbrecht *et al*., 2010), which are in the genetic background of maize inbred line W22. Here we report two insertion lines impacting the single gene encoding ATG10 (Zm00001d025765). ATG10 is an E2-like conjugating enzyme mediating the formation of ATG5-ATG12 conjugates, which is required by the ATG8 lipidation process (Mizushima *et al*., 1998; Ichimura *et al*., 2000). *atg10-Mu* (mu1052008) carries an insertion in the first exon of *ATG10*, and *atg10-Ds* (I.S06.1765) carries an insertion at the splice site between the first intron and the second exon (**Figure 3A**). RT-PCR showed that the two lines expressed no full-length transcript, although a truncated *ATG10* transcript starting from the third exon to the last exon was observed; the abundance of this truncated transcript was reduced when compared to WT W22 plants (**Figure 3B**).

**Figure 3.**
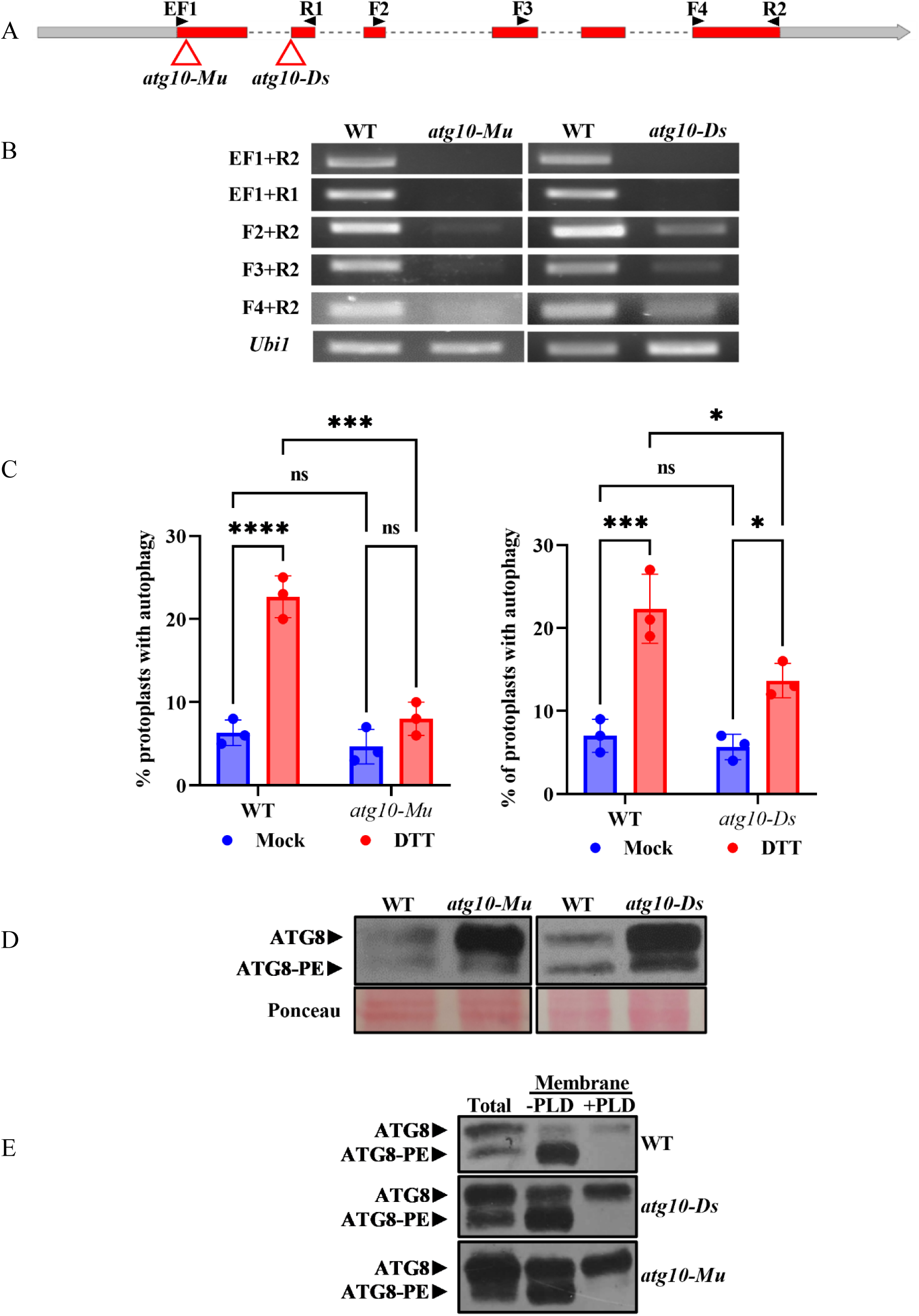
Transposon mutants in *ATG10* are defective in autophagy. **A.** Gene diagram showing the maize *ATG10* locus. Grey and red boxes represent untranslated regions and coding regions respectively. Dashed lines represent introns. Red triangles indicate the position of insertions. **B.** RT-PCR analysis of *ATG10* transcripts in *atg10-Mu* and *atg10-Ds*. RNA was extracted from field-grown wild-type W22 or homozygous mutant seedlings. 1 µg total RNA of each sample was used to synthesize cDNA, followed by PCR using specific primers to amplify *ATG10*. Locations of the primers are indicated in A. *Ubi1* was used as a control. **C.** Protoplasts were isolated from soil-grown 8- to 10-day-old wild-type W22 seedlings or homozygous *atg10-Mu* or *atg10-Ds* mutant seedlings. Protoplasts were transformed with GFP-ATG8e and incubated overnight, then treated with water as mock or 2 mM DTT for 6 h. Autophagosomes were visualized and quantified using fluorescence microscopy, and the percentage of protoplasts with activated autophagy was calculated. Asterisks indicate statistically significant differences from Tukey’s multiple comparison; *p<0.05, ***p<0.001, ****p<0.0001, ns-not significant. Error bars represent means±SD for 3 biological replicates. **D.** Proteins were extracted from the roots of 2-week-old wild-type W22 or homozygous *atg10-Mu* and *atg10-Ds* mutant seedlings. 20 µg total protein of each sample was separated by SDS-PAGE using 15% polyacrylamide gels with 6 M urea in the resolving gel and analyzed by immunoblotting with anti-ATG8 antibody. Ponceau staining was used to indicate equal loading. A representative picture is shown; 3 biological replicates showed similar results. **E.** Total protein extracts from W22, *atg10-Mu* and *atg10-Ds* were separated into soluble and membrane fractions by ultracentrifugation. The membrane fraction was solubilized with extraction buffer plus Triton X-100 and incubated with or without phospholipase D (PLD) at 37°C. Protein samples were separated by SDS-PAGE with 6M urea in the resolving gel and probed with anti-ATG8 antibody. The positions of ATG8 and ATG8-PE are indicated.

To test whether autophagy is altered in the two mutants, the autophagosome marker GFP-ZmATG8e was transiently expressed in leaf protoplasts isolated from WT W22, *atg10-Mu*, and *atg10-Ds* plants. Protoplasts were treated with or without 2 mM DTT for 6-8 hours before visualization with fluorescence microscopy. Consistent with previous results (**Figure 1E**), autophagy was significantly upregulated in WT protoplasts after DTT treatment (**Figure 3C**). However, upon DTT treatment, autophagy remained at a significantly lower level in *atg10-Mu* and *atg10-Ds* protoplasts when compared to the autophagy in WT protoplasts (**Figure 3C**). The activation of autophagy by DTT was almost completely blocked in *atg10-Mu*, while a reduced induction was observed in *atg10-Ds* (**Figure 3C**). These results suggest that autophagy is deficient in *atg10-Mu* and *atg10-Ds*, and that *atg10-Mu* may represent a stronger mutant allele. To further confirm that autophagy is deficient in these mutants, roots from 2-week-old plants grown in soil under normal conditions were collected for protein extraction, and the levels and lipidation of ATG8 was measured. ATG8 is located on autophagosomes and is degraded during autophagy (Chung *et al*., 2010). ATG8 proteins therefore accumulate in various *atg* mutants in Arabidopsis due to loss of degradation (Chung *et al*., 2010). As shown in **Figure 3D**, ATG8 proteins hyperaccumulated in both *atg10* mutants, further suggesting that autophagy is deficient in the two mutants.

In Arabidopsis, lipidated ATG8-PE is absent in an *atg10-1* mutant (Chung *et al*., 2010). In total protein extracts from the maize *atg10* mutants, a weak band was present that co-migrated with the lipidated ATG8 band observed in WT maize (**Figure 3D**). To confirm that this band corresponds to ATG8-PE, a membrane fraction was obtained by ultracentrifugation, dissolved in extraction buffer containing Triton X-100, and incubated with or without PLD at 37°C. Indeed, this band was enriched in the membrane fraction and disappeared when treated with PLD (**Figure 3E**), indicating that this is most likely ATG8-PE. It is not clear whether this small amount of lipidated ATG8 in the mutants is due to the presence of a small remaining amount of ATG10 that is below the level of detection, to a truncated ATG10 protein produced from a truncated transcript, or to the production of a small amount of lipidated ATG8 in the absence of ATG10 as can occur in in vitro conjugation reactions (Ichimura *et al*., 2004). However, it is clear that the ratio of ATG8-PE/ATG8 in *atg10* mutants is much lower than the ratio in WT (**Figure 3D**), suggesting that the efficiency of ATG8 lipidation is dramatically decreased in *atg10* mutants.

### *atg10* mutants do not have substantial growth defects

Autophagy supports vegetative growth in plants by promoting nitrogen utilization through protein degradation and nitrogen remobilization in older leaves (Guiboileau *et al*., 2012; Wada *et al*., 2015). Loss of autophagy therefore can lead to a defect in vegetative growth. Arabidopsis *atg5* and *atg7* mutants have a 50% reduction in rosette fresh weight (Minina *et al*., 2018), and in rice, an *atg7-1* mutant has reduced growth of both shoots and roots when compared to control plants (Wada *et al*., 2015). We hypothesized that maize *atg10* mutants may have reduced vegetative growth. We characterized *atg10-Mu* and *atg10-Ds* mutants, and also generated a *trans*-heterozygous *atg10-Mu*/*atg10-Ds* allele (named *atg10-MuDs*). To evaluate vegetative growth, segregating populations of *atg10-Mu*, *atg10-Ds* and *atg10-MuDs* were planted in the field following a randomized block design. Genotyping identified plants with two WT alleles and plants with two insertion alleles, which were used for comparisons. Vegetative growth-related traits including plant height, blade length, blade width and sheath length of the leaf that subtends the uppermost ear were measured at the flowering stage. *atg10-Mu* plants were on average 25 cm shorter than their WT siblings (**Figure 4A**). Leaf blade length and leaf sheath length were also reduced by ∼10% in *atg10-Mu* plants when compared to WT siblings (**Figures 4B and 4C**), but no difference was observed for leaf blade width (**Figure 4D**). However, we did not observe any significant differences for any of these traits in *atg10-Ds* and *atg10-MuDs* when compared to their siblings with two WT alleles (**Figure 4**). This suggests that any potential growth defects are minor and that ATG10 is not required for normal vegetative growth and development in maize.

**Figure 4.**
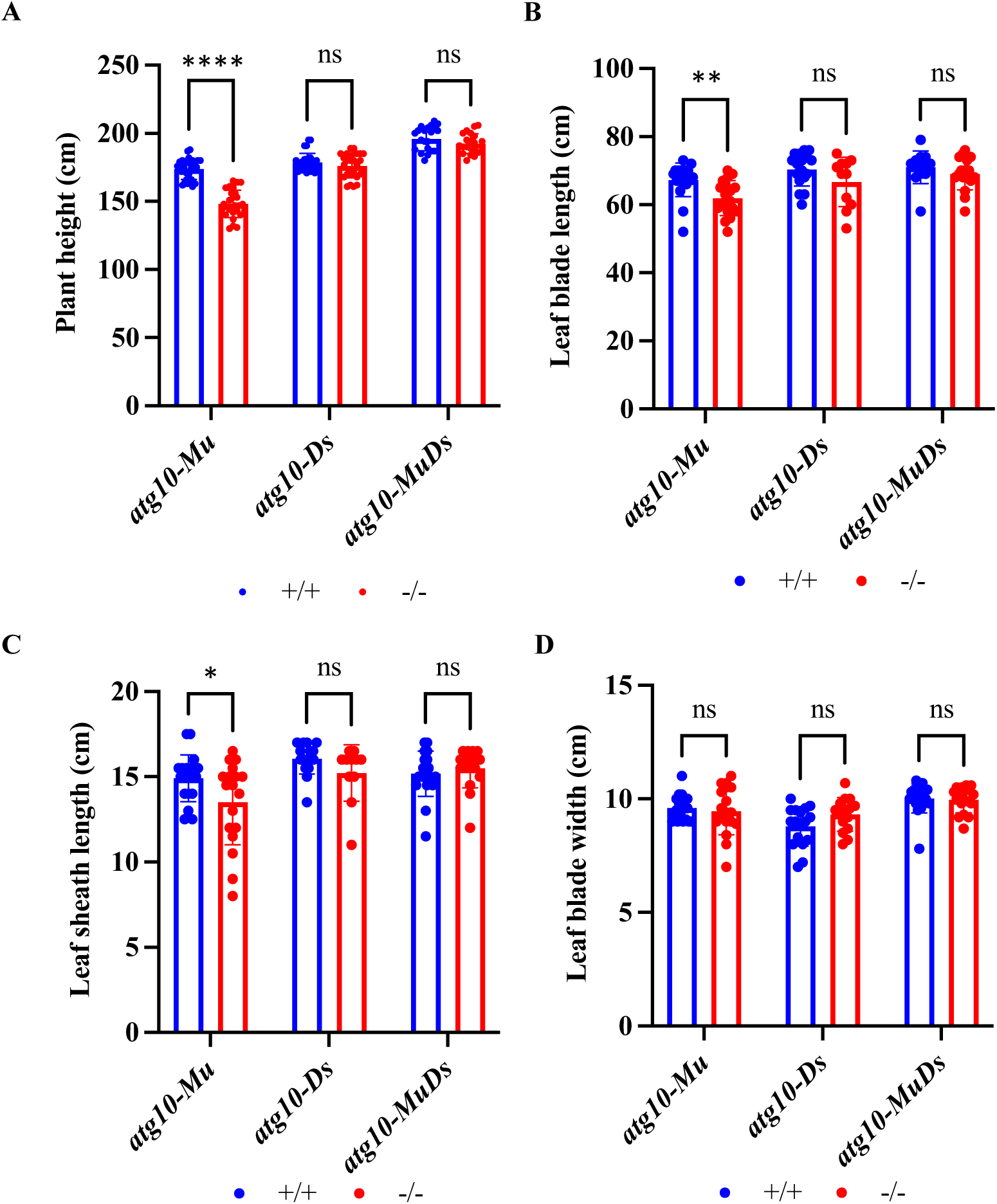
Evaluation of vegetative growth in *atg10* mutants. **A-D.** Quantification of plant height (A), leaf blade length (B), leaf sheath length (C) and leaf blade width (D). Values are means±SD (20<n<30). Dots indicate individual data points. Siblings with two wild-type alleles and siblings with two mutant alleles from segregating field-grown populations are compared. Asterisks indicate significant differences determined by Student’s t test (*p<0.05, **p<0.01, ****p<0.0001, ns-not significant).

### *bZIP60* splicing is upregulated in *atg10* mutants

We showed above that autophagy is upregulated in a *bzip60* mutant even in the absence of ER stress. A major function of autophagy during ER stress is degradation of misfolded proteins, especially larger protein aggregates (Yang *et al*., 2016). Loss of autophagy in Arabidopsis causes accumulation of ubiquitinated proteins, leading to upregulated splicing of *bZIP60* under pathogen-triggered ER stress (Munch *et al*., 2014). Thus, we hypothesized that autophagy deficiency in maize will lead to increased upregulation of the UPR upon ER stress to increase the ability to handle misfolded proteins. To test this hypothesis, maize W22, *atg10-Mu*, *atg10-Ds* and *atg10-MuDs* seeds were planted in rolls of germination papers. Plants were grown in water for 5 days and then treated with or without 2 mM DTT for 6 days. Roots were collected for RNA extraction and *bZIP60* splicing analysis. As expected, the spliced form of *bZIP60* was upregulated in DTT-treated WT roots when compared to mock-treated roots. Spliced *bZIP60* mRNA accumulated to an even higher level in DTT-treated roots of all three *atg10* genotypes (**Figure 5**), suggesting that the UPR is upregulated in *atg10* mutants subjected to ER stress, presumably as a compensatory mechanism to counteract unfolded protein accumulation.

**Figure 5.**
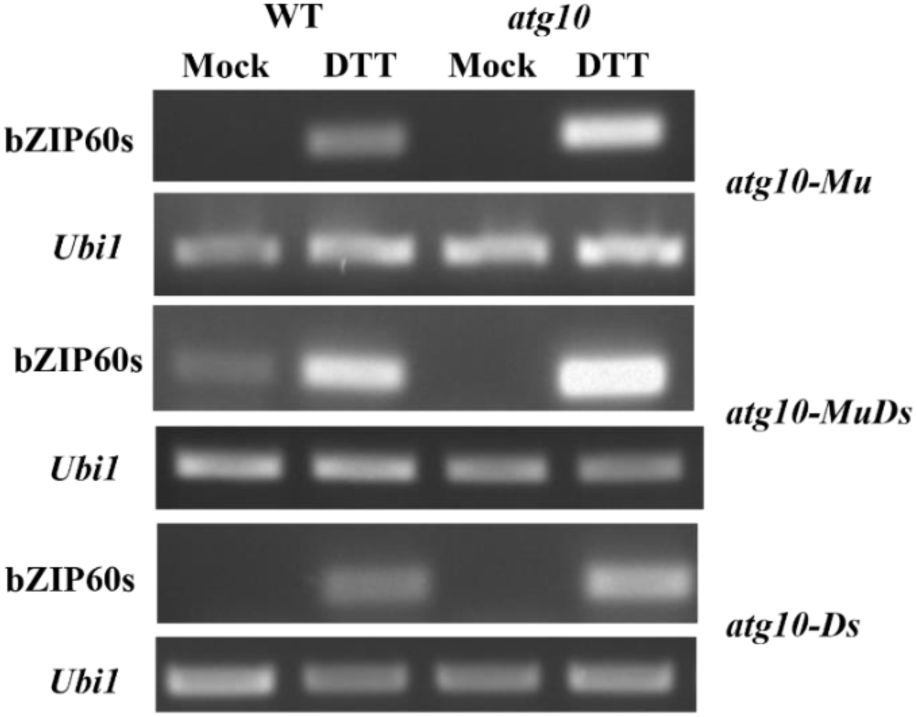
*bZIP60* splicing is increased in *atg10* mutants upon DTT treatment. WT or *atg10* seeds were planted in germination papers and grown vertically in water for 5 days, and then treated with or without 2 mM DTT for 6 days. Roots were collected for RNA extraction. The spliced and unspliced forms of *bZIP60* were amplified by RT-PCR. *Ubi1* was used as a control. A representative image is shown from three independent replicates.

### The cytoplasmic HSR, but not the UPR, is elevated in *atg10* mutants under heat stress

We have shown previously that autophagy is activated in field-grown maize during the afternoon on a hot day (Li *et al*., 2020). To determine whether this activation is due to high temperatures or to other factors, we collected leaf samples during a hot day (maximum 40°C) and a cool day (maximum 24°C) during the same week. Samples were taken from the mid part of the first fully expanded leaf every 2 hours from 11am to 5pm from multiple field-grown plants at developmental stage V7. Air temperature was recorded at the height of sample collection site (**Figure 6A**). ATG8 lipidation levels were assessed as a measure of autophagy. Autophagy was activated in W22 wild-type plants in the afternoon on the hot day, but not on the cool day, indicating that the activation was in response to the high temperature. We also measured *bZIP60* splicing in the same samples to assess activation of the UPR. Consistent with the autophagy activity, *bZIP60* splicing was also activated in the afternoon of the hot day but not the cool day.

**Figure 6.**
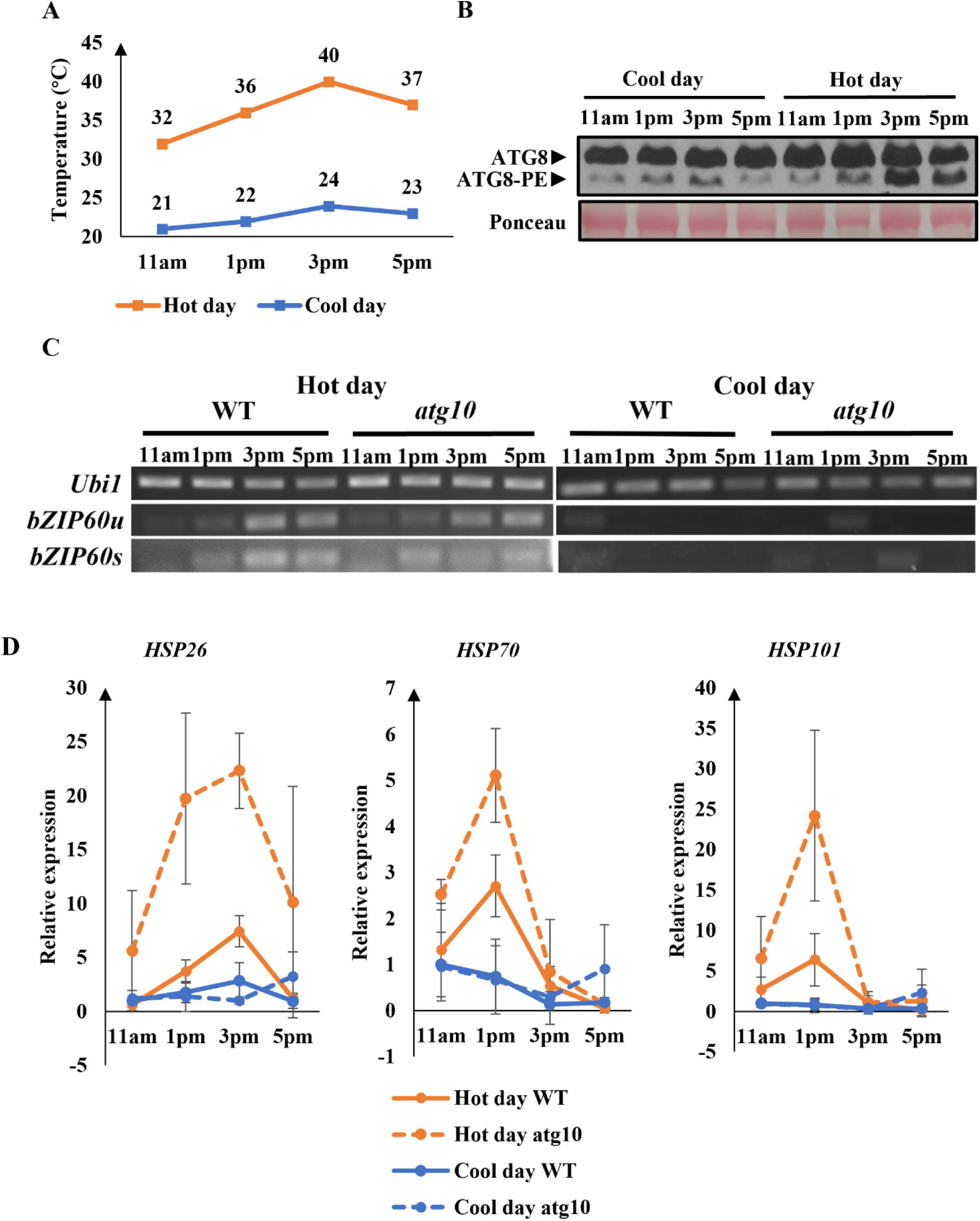
The cytoplasmic heat stress response, but not bZIP60 splicing, is increased in *atg10* mutants under heat stress in the field. Leaf samples were collected from field-grown W22 and *atg10-Mu* plants on Jun 17, 2021 (Hot day) and Jun 21, 2021 (Cool day). The first fully expanded leaf on different plants at developmental stage V7 was sampled in triplicate per genotype every 2 h (blue arrows) from 11am to 5pm. Proteins and RNA were extracted from the samples. **A.** Temperature recording on sampling days. Air temperature was measured at the height at which leaf samples were taken. **B.** ATG8 lipidation was assessed by immunoblotting to determine autophagy activity. One replicate is shown with Ponceau stained proteins as a loading control. **C.** The spliced and unspliced forms of bZIP60 were differentiated by RT-PCR. *Ubi1* was used as a control. A representative replicate is shown. **D.** RT-qPCR analysis of *HSP* gene expression. The expression level of genes at 11am in WT on the cool day was established as basal level (set to one). Values are means ± SD for three biological replicates.

As we saw that autophagy and the UPR co-occurred under field conditions in the heat, and UPR activity is increased in *atg10* mutants under ER stress when compared with wild-type plants, we hypothesized that the UPR would also be upregulated in *atg10* mutants in the field under heat stress. To test this, segregating populations of *atg10-Mu* mutants were planted in the field following a randomized block design. Plants were genotyped to identify segregants with two WT alleles and segregants with two mutant alleles, which were used for sample collection on the same days and at the same times as for W22 plants described above. bZIP60 splicing was then measured. On the hot day, but not the cool day *bZIP60s* started to increase at 1pm in all mutant genotypes and stayed at a higher level at 3pm and 5pm (**Figure 6**). However, no obvious difference was observed between WT and *atg10* mutants (**Figure 6**), indicating that the loss of autophagy has no effect on UPR activation by heat, unlike its activation by DTT.

The cytoplasmic HSR also contributes to tolerance of heat stress (Ohama *et al*., 2017), and maize bZIP60 upregulates a key transcription factor, HSFTF13, during heat stress, which in turn upregulates HSP genes (Li *et al*., 2020). HSPs prevent and resolve protein misfolding or aggregation (Mogk and Bukau, 2017). During heat stress in plants, autophagy also functions to remove protein aggregates (Zhou *et al*., 2014). We therefore assessed the potential connection between autophagy and the cytoplasmic HSR, with the hypothesis that HSPs may be upregulated in *atg10* mutants under heat stress in the field to clear protein aggregates. The expression of selected HSP genes was determined by RT-qPCR, including HSP26 (Zm00001d028408), HSP70 (Zm00001d042922) and HSP101 (Zm00001d038806), which are upregulated in response to high temperature in maize (Li *et al*., 2020). In WT plants, the expression of the three HSP genes started to rise at 1pm on the hot day (**Figure 6C**). The expression HSP70 and HSP101 peaked at 1pm and then quickly dropped, while HSP26 peaked at 3pm and started to drop at 5pm (**Figure 6C**). In *atg10* mutants, the expression patterns of the three HSP genes over the time course was similar to those in WT. However, the expression levels at the peak of the expression (1pm and 3pm for HSP26, 1pm for HSP70 and HSP101) were substantially higher than those in WT plants, suggesting that HSP genes are upregulated in *atg10* mutants under heat conditions. The loss of autophagy therefore leads to an upregulation of the UPR under ER stress and of the cytoplasmic HSR under heat stress.

## Discussion

Environmental stresses such as heat stress can disrupt protein folding and cause the accumulation of misfolded or unfolded proteins both in the ER lumen, termed ER stress, and in the cytoplasm. The cytoplasmic heat stress response aids in protein folding within the cytosol via increased production of molecular chaperones (Mogk and Bukau, 2017; Li and Howell, 2021). Plants have evolved several pathways to survive ER stress, one being the UPR (Howell, 2013), which upregulates genes involved in protein folding and ERAD to help repair or degrade unfolded proteins (Chen *et al*., 2022). The UPR factor IRE1B also activates autophagy, which functions to degrade ER fragments and unfolded proteins (Liu *et al*., 2012; Yang *et al*., 2016). Although connections between ER stress and autophagy have been demonstrated in Arabidopsis, the connections between heat stress, ER stress and autophagy in maize are still unclear.

We have shown previously that autophagy is activated in field-grown maize plants during the afternoon, and suggested that this is due to the increasing temperature during the day (Li *et al*., 2020). Here we show that autophagy is activated in the field only on a hot day and not on a cool day, indicating that the high temperature is responsible for the activation. We hypothesized that this is due to ER stress triggering the unfolded protein response, including activation of IRE1B. In a maize *bzip60* mutant, defective in a major branch of the UPR, autophagy is activated even in the absence of imposed stress (Figure 2). One possible explanation is that compromised bZIP60 function may lead to accumulation of unfolded proteins, which activates autophagy. Alternatively, the loss of bZIP60 may cause general cellular stress that triggers autophagy. In Arabidopsis, constitutive autophagy was also observed in a *bzip60* mutant (Liu *et al*., 2012). Interestingly, in Drosophila and mammals, the loss of X-BOX BINDING PROTEIN 1 (XBP1), which is a functional homolog of bZIP60, leads to upregulation of autophagy (Arsham and Neufeld, 2009; Hetz *et al*., 2009; Vidal *et al*., 2012; Zhao *et al*., 2013), as the unspliced form of XBP1 suppresses autophagy by mediating the degradation of FoxO1, a positive regulator of autophagy (Zhao *et al*., 2013). Whether a similar mechanism exists in plants is not known.

We have identified autophagy mutants with greatly reduced *ATG10* expression that have decreased autophagy activity during stress. We hypothesized that the UPR may be upregulated in these mutants to compensate for the loss of autophagy. Consistent with this hypothesis, *bZIP60* mRNA splicing was increased in *atg10* mutants compared with wild-type plants in response to ER stress triggered by DTT treatment, although constitutive UPR activation was not seen. However, while heat stress activated the UPR, no difference was seen between *atg10* mutants and wild-type plants in field-grown plants in high temperatures, suggesting a complex relationship between the UPR and autophagy, dependent on the environmental conditions. It has been reported that, in Arabidopsis, heat induces a canonical UPR gene, *ERdJ3a*, but independent of the UPR mediated by bZIP28 and bZIP60 (Howell, 2017). The promoter of *ERdJ3a* contains multiple heat shock elements, suggesting that *ERdJ3a* can be directly regulated by heat shock transcription factors (Howell, 2017). Thus, one possible explanation for the lack of difference in *bZIP60* splicing between the WT and *atg10* mutants upon heat stress is that heat stress may directly activate UPR genes without further upregulating *bZIP60* splicing.

In contrast to bZIP60 splicing, a substantial effect on the expression of cytoplasmic heat shock proteins was seen in *atg10* mutants, indicating that the cytoplasmic HSR was activated more strongly in the mutants than in wild-type plants in the heat. Insoluble proteins accumulate in autophagy-deficient mutants under heat stress (Zhou *et al*., 2013), and it is possible that in *atg10* mutants, more unfolded proteins accumulate in the cytosol under heat stress, leading to higher expression of HSPs.

In conclusion, autophagy and the UPR are both activated by DTT-triggered ER stress or by high temperature conditions in the field or in controlled environments. In a UPR-deficient mutant, autophagy is constitutively active, presumably in an attempt to compensate, and in turn the UPR is upregulated in autophagy-deficient mutants upon ER stress. However, loss of autophagy has no effect on the UPR during heat stress, but rather the cytoplasmic heat stress response is upregulated. The interplay between autophagy, the UPR and the cytoplasmic HSR therefore depends on the distinct environmental conditions.

## Materials and methods

### Plant materials

Maize transposon insertion mutants *atg10-Mu* (mu1052008) and *bzip60-2* (mu016844) were derived from the UniformMu collection (McCarty *et al*., 2005) and were obtained from the Maize Stock Center (http://maizecoop.cropsci.uiuc.edu). Insertion mutant *atg10-Ds* (I.S06.1765) was generated by the Ac/Ds project (Vollbrecht *et al*., 2010) and was kindly provided by Dr. Erik Vollbrecht. All mutant lines were in the W22 genetic background and were backcrossed to W22 three times. A trans-heterozygous *atg10-Mu*/*atg10-Ds* allele (named *atg10-MuDs*) was generated by crossing *atg10-Mu/+* with *atg10-Ds/+*. Genes and sequences were identified based on the B73 reference genome v4.

### DTT treatment

Maize W22, *atg10-Mu*, *atg10-Ds* and *atg10-MuDs* seeds were sterilized with 6% bleach for 15 min followed by washing three times with sterile water. Germination papers (Anchor Paper Co.) were soaked in Captan 50 fungicide (2.5g/L) before planting, with three sheets of germination paper used for each roll. 10 kernels were placed on the paper, the paper rolled up, and rolls placed in a 2 L beaker. Plants were grown in a growth chamber (Percival) with 16-h light/8-h dark at 26°C/20°C. Seedlings were first grown in water for 5 days and then treated with or without 2 mM DTT for 6 days. Roots of the seedlings were collected for RNA and protein extraction.

### Construction of GFP-ZmATG8e

The ZmATG8e (Zm00001d049405) coding sequence was amplified from B73 cDNA using Platinum™ Taq DNA Polymerase High Fidelity (Invitrogen). Primers used for amplification are listed in the **Supporting Table 1**. The fragment was digested with *Bgl*II and *Xba*I, and ligated into a modified pJ4-GFP transient expression vector (Igarashi *et al*., 2001; Contento *et al*., 2005), which has a GFP tag sequence at the 5’ end of the insert, driven by a 35S promoter.

### Transient expression in maize protoplasts

Transient transformation of protoplasts was performed as described (Sheen, 1991), with minor modifications. Maize W22 plants were grown in a growth chamber (Percival) with 16-h light/8-h dark at 26°C/20°C. The second leaves from 8- to 10-day-old seedlings were cut into 5 mm strips with a fresh razor blade. Leaf strips were first immersed in the enzyme solution (0.6 M mannitol, 10 mM MES pH 5.7, 1.5% (w/v) cellulase R10, 0.3% (w/v) macerozyme R10, 1 mM CaCl_2_, 5 mM 2-mercaptoethanol, 0.1% (w/v) BSA) and exposed to vacuum for 30 min, then incubated in the dark with shaking at 40 rpm for 2 hours. After incubation, protoplasts were released by shaking at 80 rpm for 5 min, and then filtered through a 75 µm mesh. Enzyme solutions containing protoplasts were centrifuged at 100 *g* for 7 min at 4°C and the supernatant discarded. Protoplasts were resuspended in wash solution (0.6 M mannitol, 20 mM KCl, 4 mM MES pH 5.7) and pelleted at 100 *g* for 5 min at 4°C. After two washes, the protoplasts were resuspended in the wash solution and incubated on ice for at least 1 hour, then pelleted at 100 g for 5 min at 4°C. Protoplasts were resuspended in MMg solution (0.6 M mannitol, 15 mM MgCl_2_, 4 mM MES pH 5.7) to a final concentration of 1∼2X10^6^ cells/mL. For transformation, 20 µL GFP-ZmATG8e plasmid (1 µg/µL), 200 µL protoplasts and 220 µL PEG solution (40% (w/v) PEG-4000, 100 mM CaCl_2_, 0.3 M mannitol) were mixed well and incubated at room temperature for 20 min. After incubation, 800 µL incubation solution (0.6 M mannitol, 4 mM KCl, 4 mM MES pH 5.7) was added to the transformation mix and mixed well, followed by centrifuge at 100 *g* for 2min. The supernatant was removed carefully. Protoplasts were resuspended in 2 mL incubation solution and transferred to a 12-well plate for 16 hours in the dark at room temperature. After incubation, protoplasts were treated with or without 2 mM DTT for 6-8 hours before visualization with fluorescence microscopy.

### Heat stress of field-grown plants

Plants were grown at the Iowa State University Curtiss Research Farm under well-watered conditions. Three different segregating populations per mutant allele were planted in three different four-row blocks, respectively, following a randomized block design. The middle part of the first fully expanded leaf was sampled every 2 h from different plants at the same developmental stage (V7). Samples were taken from three different blocks as three replicates per genotype per time point. Leaf samples were collected on Jun 17, 2021 (Hot day) and Jun 21, 2021 (Cool day). Temperature readings were taken at the height above ground from which the leaf samples were taken. Leaves were flash-frozen in the field and stored at -80°C for RNA and protein extraction.

### RNA extraction and RT-qPCR Analyses

RNA was extracted from 0.1 g plant tissue using a RNeasy Mini Kit and treated with on-column DNase (Qiagen) according to the manufacturer’s instructions. For RT-PCR analysis, 500 ng total RNA was used for cDNA synthesis (iScript cDNA Synthesis kit, Bio-Rad). cDNA was diluted 10-fold and used as template for RT-PCR and RT-qPCR analyses. Primers used are listed in **Supplemental Table 1**. RT-qPCR was performed with StepOnePlus (Applied Biosystems) using Powertrack SYBR Green Mastermix (Applied Biosystems) according to the manufacturer’s instructions. Relative gene expression levels were calculated using the 2^-ΔΔCt^ method (Livak and Schmittgen, 2001), with maize *Ubiquitin1* (*Ubi1*) as reference, and with three biological replicates.

### Genotyping

Genome DNA extraction was done as described previously (Dietrich *et al*., 2002). Leaf tissue was ground in CTAB extraction buffer (0.1 M Tris, 0.7 M NaCl, 10 mM EDTA, 1% (w/v) CTAB 1% 2-mercaptoethanol). The lysate was incubated at 65°C for 30 min with mixing by inverting the tube every 10 min. After cooling to room temperature, the lysate was mixed with 400 µL chloroform/Iso-amyl alcohol (24:1) by vortexing for 10∼15 seconds. The mix was centrifuged at room temperature for 10 min at 10,000 *g*. The supernatant was transferred to a new tube and mixed with 2.5 volumes of 100% ethanol, then incubated at -20°C for at least one hour (or overnight). After incubation, the tube was centrifugated at room temperature for 15 min at 10,000 *g*. The supernatant was discarded, and the pellet was washed with 500 µL 70% ethanol, followed by centrifugation at room temperature for 5 min at 10,000 *g*. This step was repeated twice. After the final wash, the tube was air dried, and 50 µL ddH_2_O was added to dissolve the DNA. Genotyping PCR was done using GoTaq G2 Flexi (Promega, M7801) as previously described (McCarty *et al*., 2013). Primers used are listed in **Supplemental Table 1**.

### ATG8 Lipidation Analysis

Maize tissues were harvested and immediately frozen in liquid nitrogen. ATG8 lipidation was measured as described by (Chung *et al*., 2009) with minor modifications. Plant tissues were ground in liquid nitrogen and the powder was suspended in lysis buffer (50 mM Tris-HCl at pH 8.0, 150 mM NaCl, 1 mM phenylmethanesulfonyl fluoride, 10 mM iodoacetamide, and 1X Roche cOmplete mini protease inhibitor cocktail). The crude extract was filtered through one layer of Miracloth (EMD Millipore; cat no. 475855) and then centrifuged at 2,000 *g* at 4°C for 5 min. Fifty-microgram of each protein sample was separated by SDS-PAGE using 15% polyacrylamide gels with 6 M urea in the resolving gel, and analyzed by immunoblot using anti-ATG8 antibody (Agrisera; cat. no. AS142769). Free ATG8 and lipidated ATG8 bands were quantified by ImageJ (Schneider *et al*., 2012), following the user guide (https://imagej.nih.gov/ij/docs/guide/).

### Statistical analysis

All statistical tests used GraphPad Prism 9. Student’s t-test was used for comparison between two independent groups. One-way analysis of variance (ANOVA) with Tukey’s multiple comparisons test was used for comparing multiple groups.

## Author contributions

JT and DCB designed the research; JT and ZL performed the research; JT and DCB analyzed the data; JT and DCB wrote the article; all authors approved the article before submission.

## Acknowledgements

This work was supported by grants from the National Science Foundation (Plant Genome Research Program IOS 1444339 and MCB-2040582) to DCB. The authors thanks Stephen H. Howell for helpful suggestions and discussion.

